# A root for massive crown-of-thorns starfish outbreaks in the Pacific Ocean

**DOI:** 10.1101/2021.09.13.460188

**Authors:** Nina Yasuda, June Inoue, Michael R. Hall, Manoj R. Nair, Mehdi Adjeroud, Miguel D. Fortes, Mutsumi Nishida, Nat Tuivavalagi, Rachel Ravago-Gotanco, Zac H. Forsman, Taha Basheir Hassan Soliman, Ryo Koyanagi, Kanako Hisada, Cherie A. Motti, Noriyuki Satoh

**Affiliations:** Department of Marine Biology and Environmental Science, Faculty of Agriculture, University of Miyazaki, Miyazaki 889-2192, Japan; Center for Earth Surface System Dynamics, Atmosphere and Ocean Research Institute, University of Tokyo, Kashiwa, Chiba 277-8564, Japan; Australia Institute of Marine Science, Townsville, Queensland 4810, Australia; Aquaculture Research, Extension, Training & Technology Development, College of Micronesia Land Grant Program (NIFA, USDA), Pohnpei State 96941, Federated States of Micronesia; Université de la Nouvelle-Calédonie, UMR 9220 ENTROPIE, IRD, Université de la Réunion, CNRS, IFREMER, Perpignan, France; CRIOBE - EPHE-UPVD-CNRS & Laboratoire d’Excellence “CORAIL”, PSL Université Paris, USR 3278, Perpignan, France; University of the Philippines, Diliman Quezon City 1101, Philippines; University of the Ryukyu, Nishihara, Okinawa 903-0213, Japan; Cooperative Research & Extension Department, Career & Technical Education Center, College of Micronesia-FSM, Pohnpei State 96941, Federated States of Micronesia; Marine Science Institute, University of the Philippines, Diliman, Quezon City 1101, Philippines; Coral Conservation Genetics & Restoration, Hawai’i Institute of Marine Biology, University of Hawai’i, Mānoa, HI 96744, U.S.A.; DNA Sequencing Section, Okinawa Institute of Science and Technology Graduate University, Onna, Okinawa 904-0495, Japan; Marine Genomics Unit, Okinawa Institute of Science and Technology Graduate University, Onna, Okinawa 904-0495, Japan

**Keywords:** crown-of-thorns starfish (COTS), massive outbreaks, complete mitochondrial genome, population genetics, root of larval dispersion

## Abstract

Recurring outbreaks of crown-of-thorns starfish (COTS) severely damage healthy corals in the Western Pacific Ocean. To determine the source of outbreaking COTS larvae and their dispersal routes across the Western Pacific, complete mitochondrial genomes were sequenced from 243 individuals collected in 11 reef regions. Our results indicate that Pacific COTS comprise two major clades, an East-Central Pacific clade (ECP-C) and a Pan-Pacific clade (PP-C). The ECP-C consists of COTS from French Polynesia (FP), Fiji, Vanuatu and the Great Barrier Reef (GBR), and does not appear prone to outbreaks. In contrast, the PP-C, which repeatedly spawns outbreaks, is a large clade comprising COTS from FP, Fiji, Vanuatu, GBR, Papua New Guinea, Vietnam, the Philippines, Japan, Micronesia, and the Marshall Islands. Given the nature of Pacific Ocean currents, the vast area encompassing FP, Fiji, Vanuatu, and the GBR likely supplies larvae for repeated outbreaks, exacerbated by anthropogenic environmental changes, such as eutrophication.

## Introduction

Coral reefs are the most biodiverse marine ecosystems and because they nurture edible marine species, furnish biochemicals and novel pharmaceutical leads, provide coastal protection and employment, and contribute to regional cultures, marine managers, communities and governments are calling for their preservation (De’ath et al. 2012). However, many coral reefs are currently experiencing severe, cumulative disturbances, including coral bleaching (Hughes et al. 2017), cyclones/typhoons (Harmelin-Vivien 1994), and massive outbreaks of crown-of-thorns starfish (COTS), *Acanthaster* cf. *solaris* (previously, *Acanthaster planci*) (Birkeland and Lucas 1990; Yasuda et al. 2009; Timmers et al. 2012; Hughes et al. 2014; Yasuda 2018).

COTS are considered the major and most destructive predators of reef-building corals in the Indo-Pacific (Birkeland 1990). Although they are highly fecund (Birkeland and Lucas 1990), under normal, undisturbed conditions COTS populations remain relatively constant and their impacts on coral communities are minimal (Fabricius et al. 2010). On the other hand, recent anthropogenic activities have adversely affected the marine environment resulting in an increased discharge of nutrients (Fabricius et al. 2010) and climate change (Uthicke et al. 2013), both of which are linked to increased COTS pelagic larval duration (PLD) (Yamaguch 1973). This relatively long PLD, which can last several weeks, greatly increases the overall survival rate and may assist expansion of COTS into new habitats with comparatively homogeneous populations in widespread localities (Birkeland and Lucas 1990; Vogler et al. 2013). This extended PLD, in association with strong ocean currents, is hypothesized to cause successive secondary population outbreaks of COTS, especially in the Great Barrier Reef (GBR) of Australia, and Japan (Birkeland and Lucas 1990; Benzie and Stoddart 1992; Kenchington 1997; Yasuda, 2018), with substantial loss of coral cover, thereby diminishing the integrity and resilience of reef ecosystems (Timmers et al. 2012; Hughes et al. 2014). In the GBR, one-third of coral reef damage is attributed to COTS predation (Timmers et al. 2012). Similarly, in the Ryukyu Archipelago (RA) and temperate regions of Japan, at least two waves of chronic and successive outbreaks spanning 60 years have decimated corals. Since 2000, over 980,000 COTS have been removed from reefs of Amami Island and the Ryukyus (Nakamura et al. 2014; Yasuda 2018) (website http://www.churaumi.net/onihitode/onihitode1.html), and from 2011 well over 300,000 COTS have been collected in the GBR (website http://www.environment.gov.au/marine/gbr/case-studies/crown-of-thorns), highlighting the protracted nature and high cost of programs to maintain healthy coral reefs.

Extensive studies of COTS biology, including population genetics, have been conducted (Benzie 1992; Yasuda et al. 2009; Yasuda et al. 2015; Harrison et al. 2017; Pratchett et al. 2017). For example, genetic studies based on partial mitochondrial gene sequences revealed the geographic distributions of four COTS lineages, two in the Indian Ocean, one in the Red Sea, and one in the Pacific Ocean (Vogler et al. 2008). Studies using either genes of mitochondrial cytochrome oxidase subunit I, II and III or microsatellite locus heterozygosity, or both, have generally demonstrated a genetically homogenous pattern of *A*. cf. *solari*s in the Western Pacific (Vogler et al. 2013; Tusso et al. 2016), as well as in regions associated with western boundary currents (Yasuda et al. 2009), the Hawaiian Islands (Timmers et al. 2011), French Polynesia (Yasuda et al. 2015), and the GBR (Harrison et al. 2017). However, no study has addressed which lineage of extant Pacific COTS is the oldest, what mechanisms supported their expansion across the entire Pacific Ocean, and does COTS genetic connectivity facilitate outbreaks, especially in the Western Pacific. To answer these questions, we sequenced entire mitochondrial genomes (Inoue et al. 2020) of 243 COTS specimens collected from 11 representative localities of the Pacific and conducted molecular phylogenetic analyses.

## Methods

### Acanthaster cf. solaris

A total of 243 adult crown-of-thorns starfish were collected from 2006∼2018 at reefs in the Pacific Ocean (Fig. 1). Fifty-three specimens were collected in French Polynesia, including 13 specimens from Bora-Bora, 16 from Moorea, 9 from Raiatea, and 15 from Tahiti (Supplementary Table S1). Ten specimens were collected from Fiji, 31 from Vanuatu, and 20 from the GBR (10 each from Clack and Shell Reefs). We collected 9 specimens from Papua New Guinea, 4 from the Philippines, 10 from Vietnam, 48 from the Ryukyu Archipelago of Japan and 29 from the Kagoshima Islands of Japan (Table S1). In addition, 9 specimens from Micronesia, 8 from the Marshall Islands, and 12 from the USA (9 from Hawaii and 3 from California) were also collected. Collection sites and sample numbers are reported in Supplementary Table 1.

**Figure 1.**
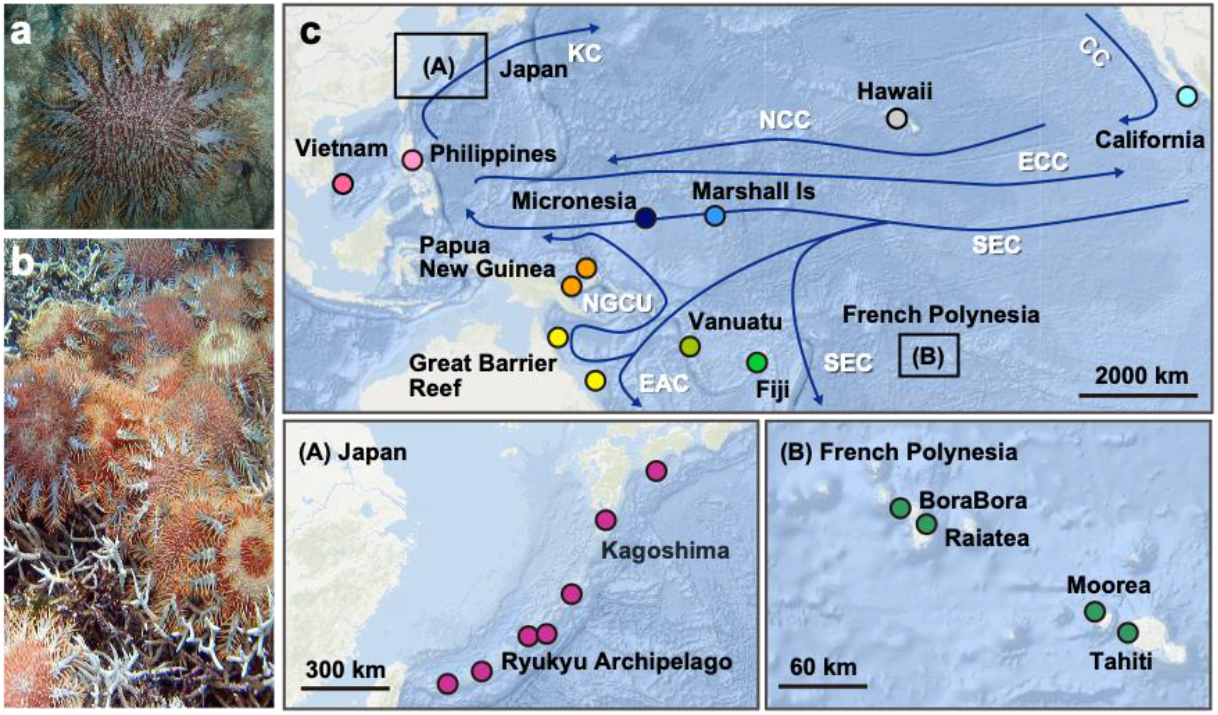
**a**, A single adult of crown-of-thorns starfish on reef-building corals. **b**, An outbreak of crown-of-thorns starfish (COTS) covering and eating scleractinian corals, causing severe damage to the reef. **c**, Collection sites of COTS in the Pacific Ocean. 243 COTS were collected at 23 locations in 14 countries, representing 11 reef regions: French Polynesia (Bora Bora, Moorea, Raiatea and Tahiti), Fiji, Vanuatu, Great Barrier Reef, Australia (GBR; Clack and Shell Reefs), Papua New Guinea (PNG), the Philippines, Vietnam, Japan, Micronesia, the Marshall Islands, and USA (Hawaii and California). Locations in Japan and French Polynesia are enlarged in (A) and (B). Red arrows show the main currents in the Pacific Ocean. KC = Kuroshio Current, CC = California Current, NEC = North Equatorial Current, ECC = Equatorial Countercurrent, SEC = South Equatorial Current, EAC = East Australian Current and NGCU = New Guinea Coastal Undercurrent.

### DNA sequencing and assembly of mitochondria genomes

Tube feet of adult COTS were dissected with scissors and fixed in 99.5% ethanol. Specimens were kept at 4°C until use for DNA sequencing. Genomic and mitochondrial DNA were extracted using the automated Nextractor® NX-48S system. Extraction was performed following the manufacturer’s protocol using an NX-48 Tissue DNA kit (Genolution Inc., Seoul, Korea). Tube foot tissue was incubated in lysis buffer overnight and extracted DNA was purified with Agencourt AMPure XP magnetic beads immediately before library preparation. DNA concentration was determined with Qubit dsDNA broad range (Thermo Scientific Inc., USA), and the quality of high molecular-weight DNA was checked using an Agilent 4150 TapeStation (Agilent, USA). PCR-free shotgun libraries were constructed using NEBNext^®^ Ultra^™^ II FS DNA Library Prep Kits for Illumina (New England BioLabs Inc, UK), following the manufacturer’s protocols. Sequencing was performed using an Illumina NovaSeq 6000 sequencer (Illumina Inc., USA).

Sequencing was performed using Illumina HiSeq 2500 and Novaseq sequencers. Approximately 10X coverage of nuclear genome DNA sequences was obtained. After removing low-quality reads, under default parameters, paired-end reads were assembled using GS *De novo* Assembler version 2.3 (Newbler, Roche) and NOVOPlasty 2.6.3 (Dierckxsens et al. 2017) with the published *A. planci* sequence l (Yasuda et al. 2006) as seed input. Usually, the largest scaffolds contained mitochondrial DNA sequences. Analysis of the genomes using MitoAnnotator (Iwasaki et al. 2013) resulted in the circular structure of the genome. That is, the genome consists of a gene set of cytochrome oxidase subunits I, II and III (COI, COII and COIII), cytochrome b (Cyt b), NADH dehydrogenase subunits 1-6 and 4L (ND1-6 and 4L), ATPase subunits 6 and 8 (ATPase6 and 8), two rRNAs, and 22 tRNAs (see Fig. 1 of 26).

As mentioned above, we collected 243 individuals representing 11 coral reef regions of the Pacific Ocean (Fig. 1) and determined the complete mitochondrial genome sequences (16,210∼16,246 bp, depending on the individual) of all specimens. Genome sequencing coverage per individual was 1,827X on average, ranging from 34X to 136,220X, indicating the data robustness from each specimen. We unambiguously aligned 16,218 bp of sequences, including 1,822 variable sites, which were used for unrooted tree analyses (Fig. 2). On the other hand, 16,219 bp of unambiguously aligned sites, including 3,159 variable sites, were used for rooted tree analyses, with mitochondrial sequences of *A. brevispinus* as an out group (Fig. 3).

**Figure 2.**
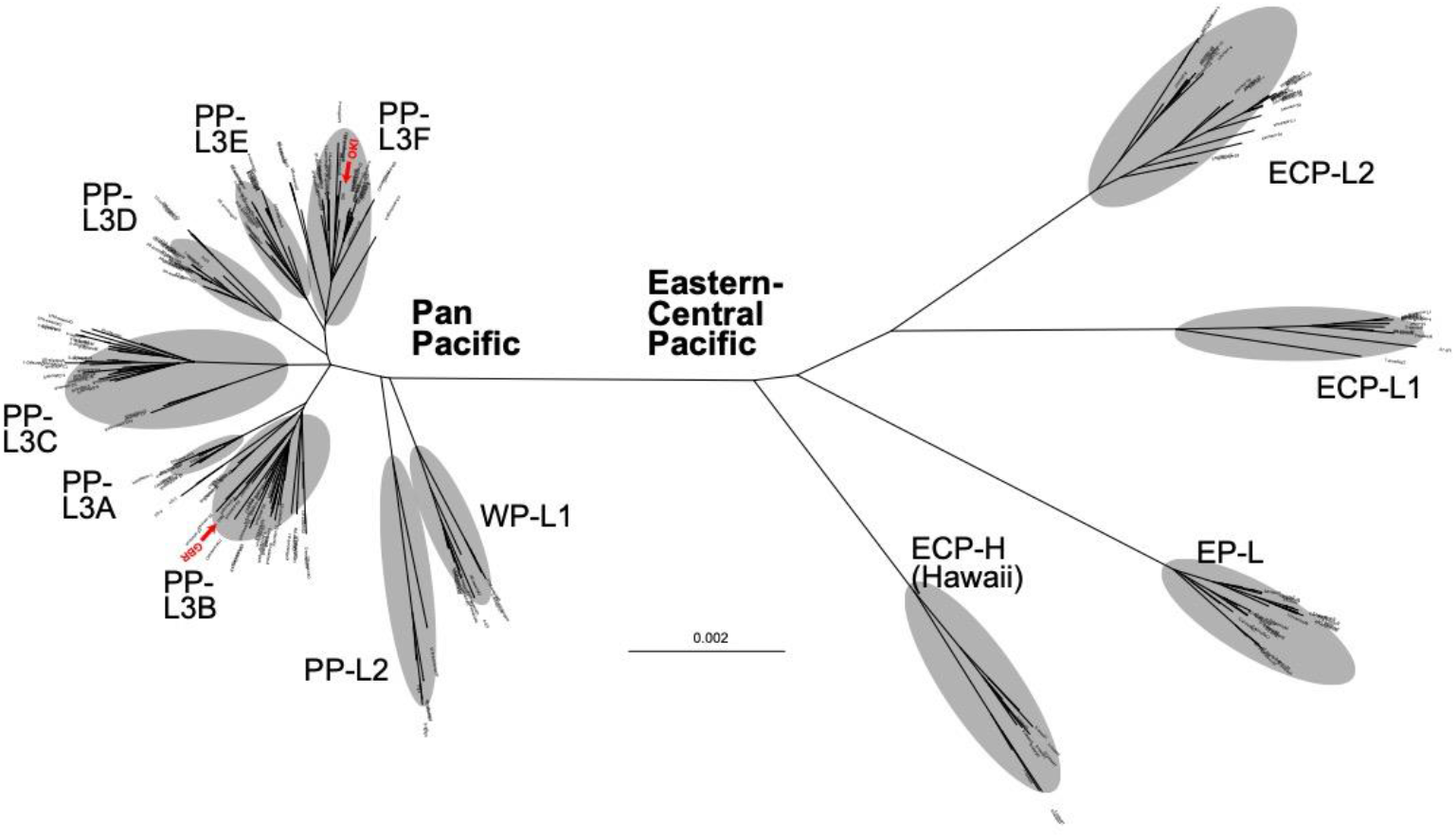
An unrooted phylogenetic tree of individual *Acanthaster* cf. *solaris* using the maximum-likelihood (ML) method, based on mitochondrial genome sequences (16,218 bp, including 1,822 variable sites). Red arrowheads indicate sequences (OKI and GBR) decoded in the genome paper (Hall et al. 2017).

**Figure 3.**
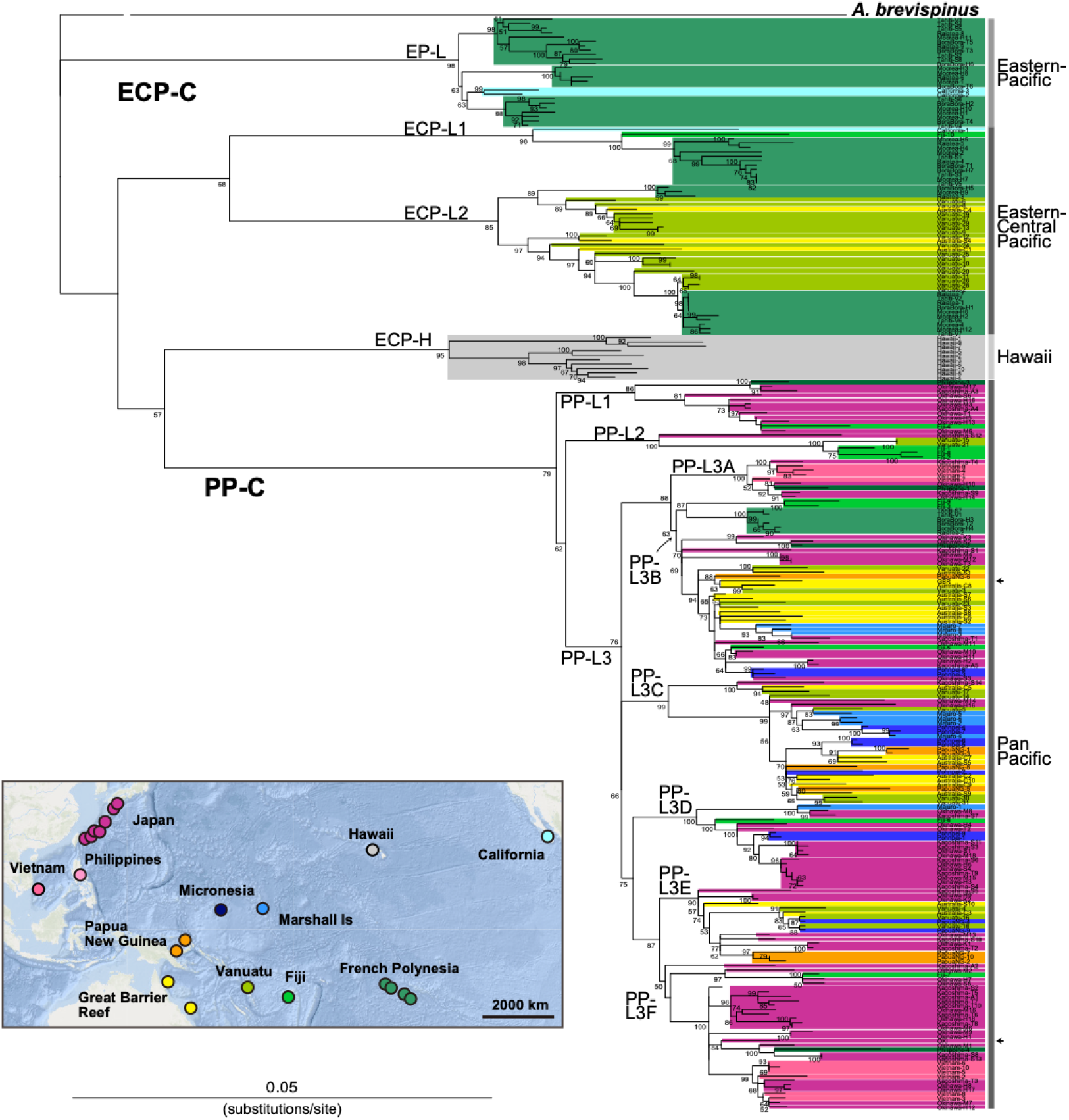
A rooted phylogenetic tree of *Acanthaster* cf. *solaris* using the maximum-likelihood (ML) method, based on mitochondrial genome sequences (16,219 bp, including 3,159 variable sites). Numbers at some nodes indicate bootstrap values (>50%) based on 100 replicates for internal branch support. Arrowheads at the right indicate sequences (OKI and GBR) decoded in the genome paper (Hall et al. 2017). The *A. brevispinus* sequence (NC_007789.1) was selected for rooting. The color relationship to sampling locations is shown in the insert (left, bottom).

### Phylogenetic analysis

Whole mitochondrial genome sequences were aligned using MAFFT (Katoh et al. 2005). Multiple sequence alignments were trimmed by removing poorly aligned regions using TRIMAL 1.2 (Capella-Gutiérrez et al. 2009) with the option “gappyout.” To examine population structures, maximum likelihood (ML) trees were created using RAxML 8.2.6 (Stamatakis 2014). Trees were estimated with the “-f a” option, which invokes rapid bootstrap analysis with 100 replicates and searches for the best-scoring ML tree, using the GTRCAT model (Stamatakis 2006).

### Principal component analysis (PCA)

Population structures were analyzed using model-free approaches. Based on mitochondrial genome sequences, principal component analysis (PCA) was performed on all individuals, using PLINK 1.9 (Purcell and Chang 2015). Pairwise genetic distances among localities were estimated with Weir and Cockerham’s *F*_ST_ (Weir and Cockerham 1984) and Nei’s genetic distance (Nei 1972) using StAMPP (Pembleton et al. 2013).

## Results and Discussion

A total of 243 adult COTS were collected from 11 representative coral reef regions (14 countries) throughout the Pacific Ocean (Fig. 1; Supplementary Table S1), including Bora Bora, Moorea, Raiatea, and Tahiti in French Polynesia, Fiji, Vanuatu, the GBR (Clack and Shell Reefs) of Australia, Papua New Guinea, the Philippines, Vietnam, Japan (the Ryukyu Archipelago and islands of Kagoshima), Micronesia, the Marshall Islands, and Hawaii and California, USA.

Complete mitochondrial genome sequences (a circular genome consisting of 16,221 bp, on average) (Inoue et al. 2010) were determined for all specimens (Supplementary Fig. S1). The mean read coverage was 1,827X, ranging from 34 to 136,220X, indicating that data were robust and suitable for establishing the complete sequence of each individual and for subsequent molecular phylogenetic analyses and principal component analysis (PCA).

An unrooted molecular phylogenetic tree was constructed for all specimens, based on 16,218 unambiguously aligned bases, including 1,822 variable sites (Fig. 2). A rooted tree using the corresponding mitochondrial sequence of *Acanthaster brevispinus* (Yasuda et al. 2006), a closely related, and possibly ancestral species of *A. planci, sensu lato* (Lucas and Jones 1976) was used as an outgroup (Fig. 3). The rooted tree was based on 16,219 unambiguously aligned bp, including 3,159 variable sites. Both trees yielded similar profiles of COTS population diversification.

Both trees indicated that COTS populations in the Pacific represent two major clades, tentatively called the East-Central Pacific clade (ECP-C) and Pan-Pacific clade (PP-C). Diversification of the two clades was evident in a long branch distance between the two in the unrooted tree. That is, the two clades are separated by 0.004 mitochondrial DNA sequence substitutions per site (Fig. 2), and there is discrete branching of the two groups in the rooted tree (Fig. 3).

The ECP-C consists of four major lineages, tentatively called the Eastern Pacific lineage (EP-L), the East-Central Pacific lineages, ECP-L1 and L2, and the Hawaiian lineage (ECP-H) (Figs. 2 and 3). The EP-L consists of COTS from French Polynesia (Tahiti, Bora Bora, Moorea, and Raiatea) and California (Fig. 3). ECP-L1, contains COTS from French Polynesia, plus populations from California and Fiji. ECP-L2 comprises two subgroups, but both include COTS of French Polynesia, Fiji, Vanuatu, and GBR (Clack and Shell Reefs). The genetic homogeneity of COTS among these French Polynesian populations was noted in a previous study (Yasuda et al. 2015). The two California COTS pertain to EPC-C, one belonging to EP-L and the other to EC1-L (Fig. 3). The external morphology of the California COTS is significantly different from counterparts in other areas of the Pacific. Specifically, they tend to have shorter arms, and were initially classified as a separate species, *Acanthaster elichii* (Timmers et al. 2012). However, allozyme analysis revealed them to have stronger affinity to COTS of the Western Pacific than to their closest geographical neighbors, the Hawaiian COTS (discussed later), and were therefore renamed *Acanthaster planci* (Nishida and Lucas 1988). This suggests a common ancestry for Eastern Pacific COTS and California COTS. Accordingly, all COTS are now classified as *Acanthaster cf solaris* (Haszprunar and Spies, 2014)

Near the root position, as viewed from the ECP-C/PP-C boundary of the unrooted tree (Fig. 2) and in the third branch of the rooted tree (Fig. 3), ten Hawaiian COTS formed a discrete group, without individuals from any other Pacific reefs (ECP-H). This genetic isolation was exceptional but had 100% bootstrap support (Fig. 3). This result agrees well with previous studies, suggesting that North Central Pacific COTS, including Hawaii, form a distinct clade among Pacific COTS (Timmers et al. 2012; Vogler et al. 2013). ECP-H is likely independent of other Pacific COTS or of cryptic COTS species. Future nuclear genomic studies should be able to confirm this possibility.

In contrast to the four lineages of ECP-C, all of which are comparatively well separated or isolated, eight lineages or subgroups of PP-C, PP-L1, PP-L2 and PP-L3A-L3F, appeared more genetically similar (Figs. 2 and 3). PP-L1, which includes COTS from Fiji, the Philippines and Japan, and PP-L2, which comprises starfish from Fiji, Vanuatu, and Japan, branched earlier and are separated from the other PP lineages (Fig. 2, Fig. 3). PP-L3 is a very large group, including not only Western Pacific COTS, but also Eastern Pacific populations from French Polynesia, Fiji, Vanuatu, GBR, Papua New Guinea, the Philippines, Japan, Micronesia, and the Marshall Islands. It consists of six lineages (PP-L3A to PP-L3F) that are not strictly geographically defined, in that each subgroup comprises individuals from several of these areas. Of special interest is PP-L3B, which has the largest geographic, including COTS from all locations of French Polynesia, Fiji, Vanuatu, GBR, Papua New Guinea, the Philippines, Vietnam, Japan, Micronesia, and the Marshall Islands. PP-L3C also includes COTS from various locations including Vanuatu, GBR, Papua New Guinea, Japan, Micronesia, and the Marshall Islands. PP-L3D includes COTS not only from Japan, Micronesia and the Marshall Islands, but also Fiji. On the other hand, PP-L3F appears to be a lineage more specific to East Asia, comprising COTS populations in the Philippines, Vietnam, and Japan.

Principle component analysis (PCA) of specimens from all sampling locations (Supplementary Table S1) supported the results of molecular phylogenetic analyses (Fig. 4). PCA resulted in five independent groups, corresponding to EP-L, ECP-L1, EPC-L2, ECP-H, and PP-L, respectively. Notably, a mixture of COTS from all locations across the Pacific was evident in PP-L (Fig. 4, upper right corner). When compared to molecular phylogeny results (Figs. 2 and 3), grouping of EP-L, ECP-L1 and EPC-H was more strongly demonstrated in PCA (Fig. 4). In addition, PCA suggested an affinity of ECP-L2 with PP-L, although this was not as strong (Fig. 4).

**Figure 4.**
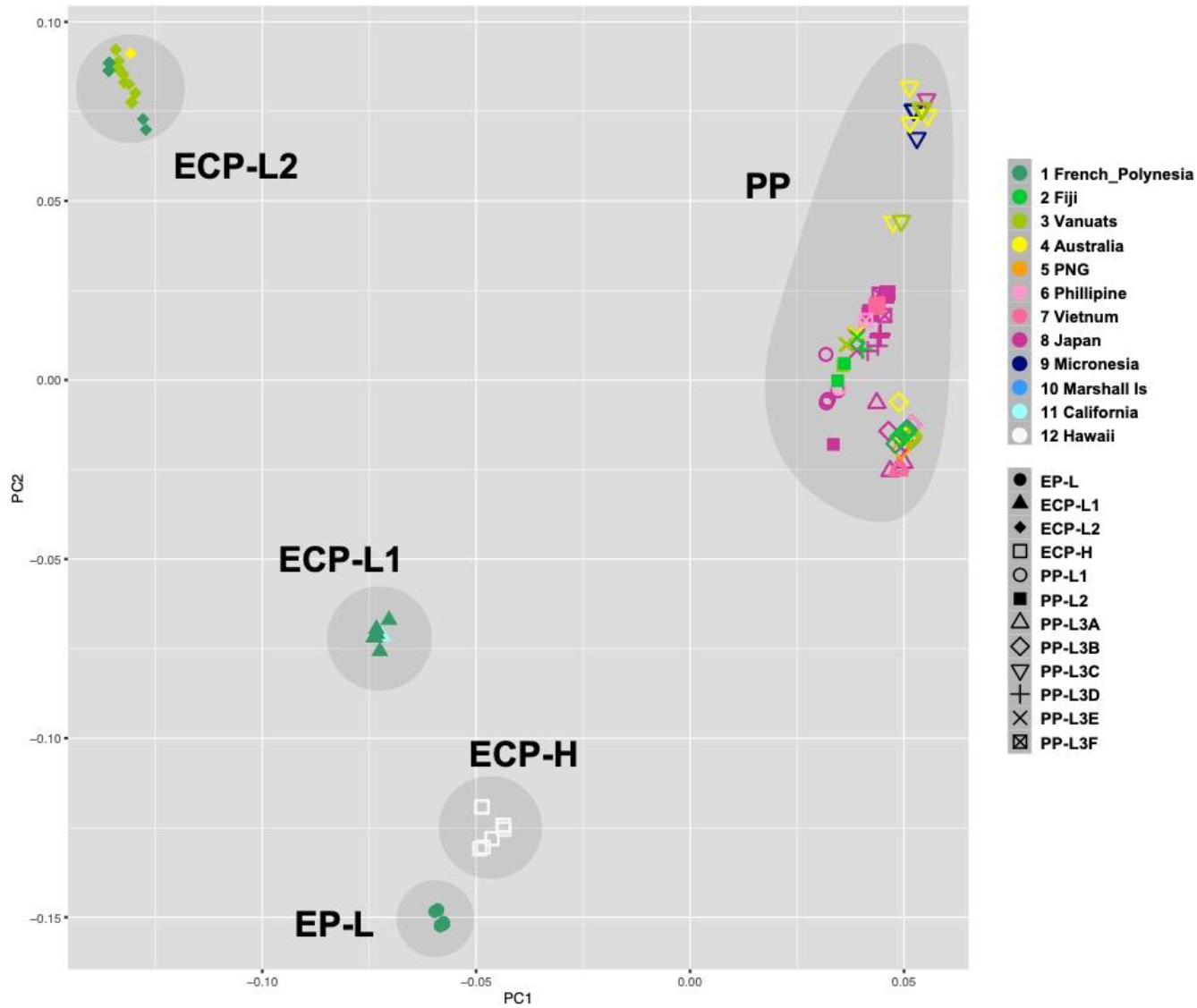
Principle Component Analysis identifies five COTS populations, EP-L, ECP-L1, ECP-L2, ECP-H, and WP-CL1/2. WP-L3 contains COTS collected from all countries, suggesting that this population is the source of repeated outbreaks in the Pacific. Color codes are shown at the right side.

The present results provide several clues regarding the evolutionary history of COTS in the Pacific Ocean. First, based on comparisons of complete mitochondrial DNA sequences, COTS in the Pacific are genetically subdivided into two major clades, ECP-C and PP-C. We speculate that because ECP-C COTS are confined to the Eastern and Central Pacific and are less affected by anthropogenic factors, they are not prone to major outbreaks, even though they show local outbreaks (Birkeland 1990). In contrast, PP-C which occurs across the entire Pacific, including more highly populated regions, spawns massive outbreaks.

ECP-C was divided into four sub-groups, EP-L, ECP-L1, ECP-L2 and ECP-H. The former three are distinguishable by their geographic distributions. EP-L is confined to four countries of French Polynesia + California, ECP-L1 encompasses French Polynesia + California + Fiji, and ECP-L2 is confined to French Polynesia, Vanuatu, and GBR. This sub-grouping suggests two possible scenarios relative to their distributional history in the Eastern and Central Pacific. One is the EP-L ancestry hypothesis, in which COTS originated in French Polynesia, experienced a bottleneck-like founder effect (Yasuda et al. 2015), and then expanded into the central and western regions, ultimately reaching the GBR. In contrast, in the ECP-L2 ancestry hypothesis, a comparatively broad region encompassing GBR, Vanuatu, Fiji and French Polynesia is the original source of COTS, from which EP-L and ECP-L1 became established as separate, independent lineages long ago. The latter scenario is the more plausible and is discussed further below.

The inclusion of Californian COTS in EP-L and ECP-L1, as well as the grouping of the independent Hawaiian lineage within EP-L, suggests that COTS larval migration in the Eastern Pacific has played an important role in their expansion across the wider Pacific. Another interesting observation is that COTS of Micronesia and the Marshall Islands may not be members of EP-L but may belong in PP-L. This suggests that the westward flow of the South Equatorial Current into the Coral Sea may become disrupted by complex topography, carrying larvae to the intersection of the Equatorial Counter Current, which is an eastward flowing, wind-driven current, thereby transporting Eastern-Central COTS larvae toward California (Wyrtki 1967) (Fig. 1). While this partially supports the ECP-L2 ancestry scenario, at present, there is no evidence to explain the origin of the Hawaiian COTS population, which arrived by unknown means and has is completely isolated. Given that Hawaiian COTS are independent of current outbreaks in the Pacific (Timmers et al. 2012), their origin remains a key question in future genomic studies.

On the other hand, PP-L contains COTS from almost all regions of the Pacific, including French Polynesia, Fiji, Vanuatu, GBR, Papua New Guinea, Vietnam, the Philippines, Japan, Micronesia, and the Marshall Islands. The two PP-L subgroups, PP-L3B and PP-L3C, both contain COTS from all these localities. It is highly likely that this type of population genetic profile reflects the trajectory of repeated outbreaks across the entire Pacific Ocean, with the exception of the U.S. population. One possible explanation is that dispersal of long-lived COTS larvae spawned in the central Pacific is facilitated by the South Equatorial Current, which flows at an average velocity of 20 nautical miles per day from Fiji and Vanuatu toward the GBR, where it bifurcates into the New Guinea Coastal Undercurrent (Treml et al. 2008; Sokolov et al. 2000). In combination with the North Equatorial Current, which originates from the Californian Current, it bifurcates into the strong Kuroshio Current that flows from the northeastern Philippines toward Japan (Qi and Lukas 1996) (Fig. 1). An earlier divergence of PP-L1 and L2, both including COTS from Fiji and Vanuatu, suggests a contribution of these COTS with western Pacific populations via the southernmost branches of the South Equatorial Current.

Further support linking repeated outbreaks to the PP-L population comes from comparisons of the entire ∼384-Mb genome sequences of the two COTS, one from the GBR and the other from Okinawa (OKI), separated by over 5,000 km (Hall et al. 2017). An unexpected result of this study was the exceptionally low heterozygosity of the genomes, 0.88% and 0.92% for the GBR and OKI populations, respectively. In addition, reciprocal BLAST analysis of scaffolds longer than 10 kb showed 98.8% nucleotide identity between the GBR and OKI genomes, evidence of the great similarity of their nuclear DNA sequences. Inclusion of these two specimens in a rooted tree (Fig. 2 and Fig. 3, arrows) revealed that GBR COTS belong to PP-L3A and Oki COTS to PP-L3F. Intriguingly, our results suggest a very strong resemblance of the nuclear genomes of these two COTS lineages.

These results raise yet another possibility with respect to the geographical extent of the distribution of various COTS lineages. Most of the COTS that belong to ECP-L1 are from Vanuatu. However, other Vanuatu COTS belong to PP-L3B, PP-L3B, or PP-L3D. Both lineages of COTS coexist in Vanuatu, one with the capacity for large outbreaks and the other without. An objective of future population genomics studies will be to sequence and compare complete genomes of both ECP-L and PP-L COTS to try to discover the genetic and genomic features that encode the capacity for outbreaks.

Based on the combined results of molecular phylogeny and PCA, it is likely that the oldest lineages of extant Pacific COTS originated from a broad Pacific region, including Fuji, Vanuatu, GBR, and French Polynesia (Fig. 5). Some populations have lived harmoniously in these regions, with some lineages moving eastward toward California. On the other hand, some COTS populations have extended their range to cover nearly all of the Pacific, and they are especially prolific in the Western Pacific (Fig. 5). The shorter branch length with highly diverse haplotypes in the admixed PP-3C implies a founder effect during their westward migration, followed by population expansion. These populations have developed a capacity for greatly enhanced larval survival, possibly triggered by anthropogenic environmental changes, such as eutrophication. Our results therefore shed light on an important issue in which regulation of future COTS outbreaks depends on a better management of this pest in the central Pacific, and better human waste management.

**Figure 5.**
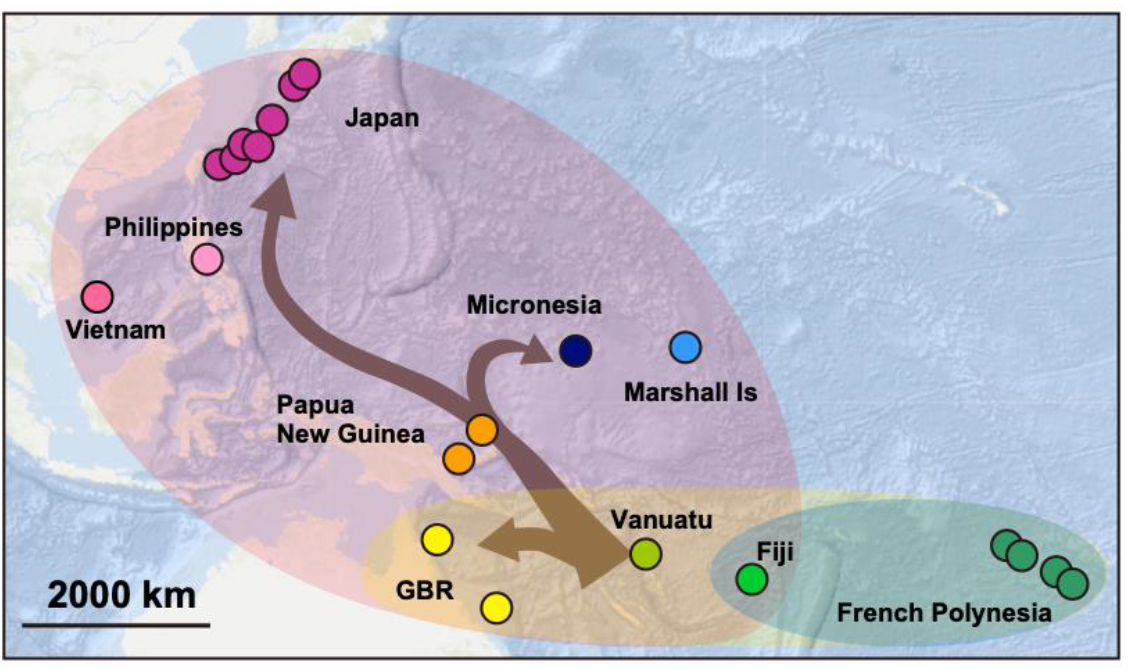
A summary diagram to show a possible root for crown-of-thorns starfish outbreaks in the Western Pacific Ocean. The Pacific hosts two major groups of COTS. The East-Central Pacific group comprises COTS from French Polynesia, Fiji, Vanuatu, and the GBR (blue and yellow). The Whole Pacific group contains COTS from the entire Western Pacific (purple). The latter has experienced repeated outbreaks, while the former has experienced local outbreaks. This suggests an importance of better management of this pest in the central Pacific region, including Fiji, Vanuatu, and the GBR.

## Data accessibility

All the sequence data are accessible under https://www.ncbi.nlm.nih.gov/bioproject/PRJDB10499.

## Authors’ contribution

N.Y., J.I., M.R.H., C.A.M. and N.S. designed the research. N.Y., M.R.H., M.R.N., M.A., M.D.F., M.N., N.T., R.R-C. and S.H.F. collected samples. T.B.H.S. and R.K. sequenced COTS mitochondrial DNA. N.Y., J.I., K.H., C.A.M. and N.S. analyzed data. N.Y., J.I., C.A.M. and N.S. wrote the manuscript with input from all authors. All authors gave final approval for publication.

## Acknowledgements

We thank the following people for their help with sample collection: Dr. Hugh Sweatman and the AIMS Bioresources Library for GBR samples, Dr. Molly Timmers for Hawaiian samples, Geoff Jones and Jeff Kinch for Papua New Guinean samples, Monal Lal for Fijian samples, Christina Shaw for Vanuatu samples, Hoang Dinh Chieu for Vietnamese samples, and Hiromitsu Ueno for Japanese samples. The DNA Sequencing Section of OIST is acknowledged for its expert help with genome sequencing.

## Figure Legends

**Figure S1**. An alignment of the complete mitochondrial DNA sequences of *Acanthaster planci* (NC_007788.1 and specimen name, M2), All sequence data are accessible at: https://www.ncbi.nlm.nih.gov/bioproject/PRJDB10499.

## References

Benzie, J., and Stoddart, J. (1992). Genetic structure of outbreaking and non-outbreaking crown-of-thorns starfish (Acanthaster planci) populations on the Great Barrier Reef. Marine Biology 112, 119–130.

Benzie, J. (1992). Review of the genetics, dispersal and recruitment of crown-of-thorns starfish (Acanthaster planci). Marine and Freshwater Research 43, 597–610.

Birkeland, C. (1990). Acanthaster planci: major management problem of coral reefs / authors, Charles E. Birkeland, John S. Lucas, (Boca Raton: CRC Press).

Birkeland, C., and Lucas, J. (1990). Acanthaster planci. Major Management Problem of Coral Reefs.

Capella-Gutiérrez, S., Silla-Martínez, J.M., and Gabaldón, T. (2009). trimAl: a tool for automated alignment trimming in large-scale phylogenetic analyses. Bioinformatics 25, 1972–1973.

De’ath, G., Fabricius, K.E., Sweatman, H., and Puotinen, M. (2012). The 27– year decline of coral cover on the Great Barrier Reef and its causes. Proceedings of the National Academy of Sciences 109, 17995–17999.

Dierckxsens, N., Mardulyn, P., and Smits, G. (2017). NOVOPlasty: de novo assembly of organelle genomes from whole genome data. Nucleic acids research 45, e18–e18.

Fabricius, K.E., Okaji, K., and De’ath, G. (2010). Three lines of evidence to link outbreaks of the crown-of-thorns seastar Acanthaster planci to the release of larval food limitation. Coral Reefs 29, 593–605.

Hall, M.R., Kocot, K.M., Baughman, K.W., Fernandez-Valverde, S.L., Gauthier, M.E., Hatleberg, W.L., Krishnan, A., McDougall, C., Motti, C.A., and Shoguchi, E. (2017). The crown-of-thorns starfish genome as a guide for biocontrol of this coral reef pest. Nature 544, 231–23436.

Harmelin-Vivien, M.L. (1994). The effects of storms and cyclones on coral reefs: a review. Journal of Coastal Research, 211-231.

Harrison, H.B., Pratchett, M.S., Messmer, V., Saenz-Agudelo, P., and Berumen, M.L. (2017). Microsatellites reveal genetic homogeneity among outbreak populations of crown-of-thorns starfish (Acanthaster cf. solaris) on Australia’s Great Barrier Reef. Diversity 9, 16.

Haszprunar, G., and Spies, M. (2014). An integrative approach to the taxonomy of the crown-of-thorns starfish species group (Asteroidea: Acanthaster): A review of names and comparison to recent molecular data. Zootaxa 25, 271–284.

Hughes, R.N., Hughes, D.J., and Smith, I.P. (2014). Limits to understanding and managing outbreaks of crown-of-thorns starfish (Acanthaster spp.). Oceanography and Marine Biology: An Annual Review 52, 133–200.

Hughes, T.P., Kerry, J.T., Álvarez-Noriega, M., Álvarez-Romero, J.G., Anderson, K.D., Baird, A.H., Babcock, R.C., Beger, M., Bellwood, D.R., and Berkelmans, R. (2017). Global warming and recurrent mass bleaching of corals. Nature 543, 373–377.

Inoue, J., Hisata, K., Yasuda, N., and Satoh, N. (2020). An Investigation into the genetic history of Japanese populations of three Starfish, Acanthaster planci, Linckia laevigata, and Asterias amurensis, based on complete mitochondrial DNA sequences. G3: Genes, Genomes, Genetics 10, 2519–2528.

Iwasaki, W., Fukunaga, T., Isagozawa, R., Yamada, K., Maeda, Y., Satoh, T.P., Sado, T., Mabuchi, K., Takeshima, H., and Miya, M. (2013). MitoFish and MitoAnnotator: a mitochondrial genome database of fish with an accurate and automatic annotation pipeline. Mol Biol Evol 30, 2531–2540.

Katoh, K., Kuma, K.-i., Toh, H., and Miyata, T. (2005). MAFFT version 5: improvement in accuracy of multiple sequence alignment. Nucleic acids research 33, 511–518.

Kenchington, R. (1977). Growth and recruitment of Acanthaster planci (L.) on the Great Barrier Reef. Biol Conserv 11, 103–118.

Lucas, J.S., and Jones, M.M. (1976). Hybrid crown-of-thorns starfish (Acanthaster planci x A. brevispinus) reared to maturity in the laboratory. Nature 263, 409–412.

Lucas, J.S. (1982). Quantitative studies of feeding and nutrition during larval development of the coral reef asteroid Acanthaster planci (L.). Journal of Experimental Marine Biology and Ecology 65, 173–193.

Nakamura, M., Okaji, K., Higa, Y., Yamakawa, E., and Mitarai, S. (2014). Spatial and temporal population dynamics of the crown-of-thorns starfish, Acanthaster planci, over a 24-year period along the central west coast of Okinawa Island, Japan. Marine Biology 161, 2521–2530.

Nei, M. (1972). Genetic distance between populations. The American Naturalist 106, 283–292.

Nishida, M., & Lucas, J. S. (1988). Genetic differences between geographic populations of the crown-of-thorns starfish throughout the Pacific region. Marine Biology, 98(3), 359–368.

Pembleton, L.W., Cogan, N.O., and Forster, J.W. (2013). St AMPP: An R package for calculation of genetic differentiation and structure of mixed-ploidy level populations. Molecular ecology resources 13, 946–952.

Pratchett, M.S., Dworjanyn, S., Mos, B., Caballes, C.F., Thompson, C.A., and Blowes, S. (2017). Larval survivorship and settlement of crown-of-thorns starfish (Acanthaster cf. solaris) at varying algal cell densities. diversity 9, 2.

Pratchett, M.S., Caballes, C.F., Wilmes, J.C., Matthews, S., Mellin, C., Sweatman, H., Nadler, L.E., Brodie, J., Thompson, C.A., and Hoey, J. (2017). Thirty years of research on crown-of-thorns starfish (1986–2016): scientific advances and emerging opportunities. Diversity 9, 41.

Purcell, S., and Chang, C. (2015). PLINK 1.9. Available from: www.cog-genomics.org/plink/1.9.

Qiu, B., and Lukas, R. (1996). Seasonal and interannual variability of the North Equatorial Current, the Mindanao Current, and the Kuroshio along the Pacific western boundary. Journal of Geophysical Research: Oceans 101, 12315–12330.

Sokolov, S., and Rintoul, S. (2000). Circulation and water masses of the southwest Pacific: WOCE section P11, Papua New Guinea to Tasmania. Journal of marine research 58, 223–268.

Stamatakis, A. (2006). RAxML-VI-HPC: maximum likelihood-based phylogenetic analyses with thousands of taxa and mixed models. Bioinformatics 22, 2688–2690.

Stamatakis, A. (2014). RAxML version 8: a tool for phylogenetic analysis and post-analysis of large phylogenies. Bioinformatics 30, 1312–1313.

Timmers, M.A., Andrews, K.R., Bird, C.E., deMaintenton, M.J., Brainard, R.E., and Toonen, R.J. (2011). Widespread dispersal of the Crown-of-Thorns sea star, Acanthaster planci, across the Hawaiian Archipelago and Johnston Atoll. Journal of Marine Biology 2011, 10.

Timmers, M., Bird, C., Skillings, D., Smouse, P., and Toonen, R. (2012). There’s No Place Like Home: Crown-of-Thorns Outbreaks in the Central Pacific Are Regionally Derived and Independent Events. PLoS ONE 7, e31159.

Treml, E.A., Halpin, P.N., Urban, D.L., and Pratson, L.F. (2008). Modeling population connectivity by ocean currents, a graph-theoretic approach for marine conservation. Landscape Ecology 23, 19–36.

Tusso, S., Morcinek, K., Vogler, C., Schupp, P.J., Caballes, C.F., Vargas, S., and Wörheide, G. (2016). Genetic structure of the crown-of-thorns seastar in the Pacific Ocean, with focus on Guam. PeerJ 4, e1970.

Uthicke, S., Pecorino, D., Albright, R., Negri, A.P., Cantin, N., Liddy, M., Dworjanyn, S., Kamya, P., Byrne, M., and Lamare, M. (2013). Impacts of ocean acidification on early life-history stages and settlement of the coral-eating sea star Acanthaster planci. PLoS ONE 8, e82938.

Vogler, C., Benzie, J., Lessios, H., Barber, P., and Worheide, G. (2008). A threat to coral reefs multiplied? Four species of crown-of-thorns starfish. Biol Lett 4, 696–699.

Vogler, C., Benzie, J.A.H., Tenggardjaja, K., Ambariyanto, Barber P.H., and Wörheide, G. (2013). Phylogeography of the crown-of-thorns starfish: genetic structure within the Pacific species. Coral Reefs 32, 515–525.

Weir, B.S., and Cockerham, C.C. (1984). Estimating F-statistics for the analysis of population structure. evolution, 1358–1370.

Wyrtki, K. (1967). Equatorial Pacific Ocean1. Int J Oceanol & Limnol 1, 117–147.

Yamaguchi, M. (1973). Early life history of coral reef asteroids, with special reference to Acanthaster planci (L.). Biology and Geology of Coral Reefs Biology 1 II, 369–389.

Yasuda, N., Hamaguchi, M., Sasaki, M., Nagai, S., Saba, M., & Nadaoka, K. (2006). Complete mitochondrial genome sequences for Crown-of-thorns starfish Acanthaster planci and Acanthaster brevispinus. BMC genomics, 7(1), 17.

Yasuda, N., Nagai, S., Hamaguchi, M., Okaji, K., Gérard, K., and Nadaoka, K. (2009). Gene flow of Acanthaster planci (L.) in relation to ocean currents revealed by microsatellite analysis. Molecular Ecology 18, 1574–1590.

Yasuda, N., Taquet, C., Nagai, S., Yoshida, T., and Adjeroud, M. (2014). Genetic connectivity of the coral-eating sea star Acanthaster planci during the severe outbreak of 2006–2009 in the Society Islands, French Polynesia. Marine Ecology, n/a-n/a.

Yasuda, N. (2018). Distribution Expansion and Historical Population Outbreak Patterns of Crown-of-Thorns Starfish, Acanthaster planci sensu lato, in Japan from 1912 to 2015. In Coral Reef Studies of Japan, A. Iguchi and C. Hongo, eds. (Singapore: Springer Singapore), pp. 125–148.

